# Pharmacological blockade of glutamatergic input to the lateral habenula modulates consumption of palatable diet components in male Wistar rats

**DOI:** 10.1101/2024.09.02.610523

**Authors:** Margo Slomp, Milou T. Spitters, Jolinde L. van Bergen, Astrid A.S. van Irsen, Tess Kool, Leslie Eggels, Joram D. Mul, Susanne E. la Fleur

## Abstract

The lateral habenula (LHb), a small epithalamic nucleus, modifies downstream midbrain dopamine neuron output to regulate negative state and aversion. Furthermore, specific glutamatergic input, from, among others, the lateral hypothalamus and central amygdala to LHb modulates consumption of (palatable) diet components. However, it is currently unclear if blockade of all glutamatergic input to the LHb is sufficient to alter eating behavior.

Here, we used a pharmacological approach to inhibit all glutamatergic input to the LHb by bilateral infusion of either an AMPA/kainate receptor antagonist (CNQX) or an NMDA receptor antagonist (AP5) in the LHb of male Wistars rats. We then measured consumption of various palatable diets a control diet, a free-choice high-fat diet (fcHFD), a free-choice high-sugar diet (fcHSD), and a free-choice high-fat high-sugar diet (fcHFHSD)] at various timepoints up to 24h following infusion. Rats consumed their respective diets for 14 days before infusion of vehicle, CNQX or AP5, performed in counter-balanced random order.

Infusion of CNQX or AP5 did not acutely (*i*.*e*. 1, 3, or 6h following infusion) affect consumption of a fcHFHSD component. Infusion of AP5 decreased fat intake at later time points (*i*.e. 10 or 24h following infusion) in fcHFHSD- and fcHFD-fed, but not fcHSD-fed, rats. Combined infusion of CNQX and AP5 decreased sucrose water consumption at 24h following infusion in fcHFHSD-fed rats. Collectively, these observations indicate that blocking glutamatergic transmission in the LHb does not have a major impact on acute consumption of palatable free-choice diet components. Nonetheless, more subtle long-term effects were observed, suggesting a modulatory role of LHb in eating behavior in the current experimental set-up.

## Introduction

The habenula is a small, evolutionary well-conserved structure in the brain [1-3], which is subdivided into a lateral (LHb) and medial (MHb) part [4]. These subdivisions have distinct gene expression profiles and anatomical connectivity, resembling their different functions in reward processing and mood regulation [5-10]. The LHb predominantly contains glutamatergic neurons and relatively few GABAergic neurons [6, 11] and exhibits spontaneous activity in a circadian manner, with increased activity during the (subjective) day [12-14]. The LHb is considered a hub between various limbic structures and midbrain neuromodulator systems which signals and processes negative state and aversion [15, 16]. The LHb regulates dopamine function through direct, glutamatergic projections to dopaminergic neurons within the ventral tegmental area (VTA) [17]. Via this projection, the LHb can induce the release of dopamine, but the LHb also sends glutamatergic afferents to GABAergic neurons of the rostromedial tegmental nucleus (RMTg) [18], which then in turn can inhibit VTA dopaminergic neurons [19]. These direct and indirect projections allow the LHb to bi-directionally modulate dopamine signaling, and thus modulate goal-directed and motivational behaviors [4, 18-22]. This modulation of reward-seeking behavior likely also extends to eating behavior of palatable diets.

The LHb receives and integrates information from a wide range of brain regions, including entopeduncular nucleus (EPN) [23-26], lateral hypothalamus (LH) [27-29], basal forebrain [30, 31], and reciprocal connections from the VTA [32, 33], to guide behavior. Several of these inputs co-release GABA and glutamate, and the balance between these two crucial neurotransmitters is thought to maintain normal activity of the LHb, as this region contains relatively few GABAergic interneurons [27, 34, 35]. Notably, a disbalance in the GABA::glutamate ratio has been implicated in the regulation of mood and behavior [27, 34-36]. Indeed, LHb hyperactivity has been observed in depressed humans and chronically stressed animals [37-46]. Depression-related LHb hyperactivity is suggested to be driven by AMPA receptor-mediated potentiation [47], although NMDA antagonists have also been reported to decrease depression-related symptoms [48, 49]. Recently, consumption of a high-fat diet (HFD) has also been linked to increased LHb activity [50].

Thus, palatable diet consumption is likely to modulate LHb excitability, and in turn, the LHb could regulate food-seeking behavior in a similar manner as other rewards. Indeed, glutamatergic innervation of the LHb originating from the LH negatively regulates the consumption of a highly caloric solution [29]. This LH->LHb connection has been studied extensively and is likely linked to aversive taste processing [51, 52], and also seems modulated by satiety state and energy metabolism-related hormones like leptin and ghrelin [52, 53]. Another glutamatergic input to the LHb, from the basal forebrain, limits consummatory behavior of a palatable diet during a fasted state [31]. Furthermore, activation of µ-opioid signaling in the LHb increased intake of a standard (chow) diet, whereas it decreased consumption of sweetened fat [54]. Neuropeptide Y (NPY) signaling within the habenula also regulates consummatory behavior of a palatable diet [50], as inhibition of NPY receptor 1 (NPY1R) in the LHb increased HFD intake; however, when eating of an HFD was combined with stress, LHb-specific knockdown of NPY1R actually decreased food intake [50]. This suggests a stress-dependent role of LHb NPY signaling in the control of palatable feeding. Notably, one possible LHb NPY input important for stress-related feeding is a glutamatergic projection from the central amygdala [50, 55].

Collectively, the literature suggests a potential role for mostly glutamatergic inputs to the LHb in the regulation of palatable diet consumption. To study this process, various diets are available with pellets containing either high levels of fat, sugar, or both. However, the lack of choice in these diet components as well as the respective timing of intake of fat or sugar makes it far from ideally suited to model the etiology of human obesity [56]. Therefore, our lab commonly uses a four-component free-choice high-fat high-sugar diet (fcHFHSD), consisting of prefabricated nutritionally complete pellets (chow), a dish of beef tallow, a bottle with 30% sucrose solution, and a bottle of tap water, which are all freely as well as simultaneously available [56-58]. Consumption of fat and sugar can potentiate reward via additive mechanisms which have their effect on brain function, energy metabolism, and subsequent consummatory behavior [59-62]. Therefore, we hypothesized that blocking all glutamatergic input to the LHb would modulate consumption of palatable diet components of the fcHFHSD, or the three-component free-choice high-fat diet (fcHFD) or three-component free-choice high-sugar diet (fcHSD), compared to a standard no-choice chow diet. In this study, we tested this by inhibiting glutamatergic input to the LHb using a pharmacological approach [*i*.*e*. by infusing either an AMPA/kainate (CNQX) or an NMDA (AP5) receptor antagonist] and measuring consumption of individual diet components up to 24h.

## Methods

### Animals

All experiments were performed in male Wistar rats (Charles River Breeding Laboratories, Sulzfeld, Germany), with initial weights ranging between 240-280 grams. For the duration of the experiment, animals were held in a temperature-(21 ± 2°C), humidity-(60 ± 5%), and light-controlled (12:12hr light/dark; lights on 07:00-19:00) room with background noise (radio). All animals had *ad libitum* access to a standard diet (Teklad global diet 2918; 24% protein, 58% carbohydrate, and 18% fat, 3,1 kcal/g, Envigo, Horst, The Netherlands) and tap water and were group-housed during a minimum of seven days of acclimatization. All experiments were approved by the animal ethics committee of the Royal Dutch Academy of Arts and Sciences (KNAW, Amsterdam, the Netherlands) and in accordance with the guidelines on animal experimentation of the Netherlands Institute for Neuroscience and Dutch legal ethical guidelines.

### Stereotaxic surgery

Rats were anesthetized with an intraperitoneal injection of a mix of 80 mg/kg Ketamine (Eurovet Animal Health, Bladel, the Netherlands), 8 mg/kg Rompun® (xylazine, Bayer Health Care, Mijdrecht, the Netherlands) and 0.1 mg/kg Atropine (Pharmachemie B.V., Haarlem, the Netherlands). Lidocaine (Xylocaine 10% spray) was applied to the periosteum to provide additional local analgesia. Animals were placed on a heating pad in the stereotaxic apparatus (Kopf®, David Kopf Instruments, Tujunga, California). Two 26-gauge stainless steel guide cannula (C315G-SPC 5▫mm, Plastics One, Bilaney Consultants GmbH, Düsseldorf, Germany) aimed bilaterally at the LHb were implanted with a 10° angle at anteroposterior (AP) -3.7, mediolateral (ML) ±1.7, dorsoventral (DV) -4.6 (from the skull) according to the Rat Paxinos atlas (Paxinos & Charles Watson, 2007). Cannulas were secured to the skull with 4 screws and dental cement. Rats received Carprofen (5 mg/kg BW, subcutaneous) during surgery and the first post-surgery day. During the recovery period of 7 days, food and water intake and body weight were measured daily. From surgery onwards, rats were individually housed with a gnawing stick as cage enrichment.

### Diet exposure

After recovery and when pre-operative bodyweight was again reached, typically after seven days, the experiment continued. Rats first received a bilateral infusion of 0.5µl phosphate-buffered saline (PBS) to familiarize the animals with the procedure of infusions. Rats were distributed over two experiments: in one experiment the animals received the glutamatergic AMPA/Kainate receptor antagonist CNQX, and in the other experiment animals received DL-AP5, a glutamatergic NMDA receptor antagonist. In each experiment, animals were randomly divided over four different diet groups: a chow-fed group, a fcHFHSD group, a fcHFD group and a fcHSD group. All groups had access to chow and water *ad libitum* and the fcHFD, fcHSD and fcHFHSD groups had access to a dish of saturated fat (beef tallow, Vandemoortele, Belgium), a 30% sucrose (table sugar) solution, or both, respectively.

### Infusions

After two weeks of diet exposure to their respective diet, rats received infusions of either phosphate-buffered saline, disodium-CNQX (0,5 µl 1,8mM, Abcam; ab120044)[63] or DL-AP5 (0,5 µl 80mM, Abcam; ab120271)[48] in a randomized, counterbalanced within-animal design. Infusions were performed twice a week, and animals were maintained on their respective diet throughout the experiment. On infusion days, food items (but not water) were removed at Zeitgeber Time (ZT) 0 (lights off) and 5 grams of chow was provided to have a controlled level of hungriness. Then, around 3 hours after lights-off (ZT15), infusions were performed in the dark. Animals always consumed the 5 grams of chow provided. Infusions were performed using an injector connected to a 10µL Hamilton syringe placed in an infusion pump (Harvard Apparatus) with a rate of 0.3 µL/min while rats were gently held. The injectors were left in the guide cannula for 1 min after completion of the infusion to allow for diffusion. Pre-weighed amounts of all food components were provided and measured after various time points (1h, 3h, 6h, 10h, 24h, depending on the experiment). CNQX and AP5 infusions were performed in separate groups of animals.

### Tissue collection and cannula placement

Two to five days after the final infusion, the animals were anesthetized by O2/CO2 followed by rapid decapitation. To assess body composition, the following fat pads were dissected and weighted immediately (unilaterally): mesenteric (MWAT), epididymal (EWAT), perineal (PWAT), and subcutaneous (SWAT) white adipose tissue. Brains were dissected, freshly frozen on dry ice, and stored at -80°C for cannula placement determination. Coronal brain slices of 35 µm were cut on a cryostat (CryoStar NX50, Thermo Scientific) and were collected on Superfrost ++ slides. These sections were fixed with a 4% paraformaldehyde solution and stained for Nissl using thionine. Stained sections were examined using a microscope (Leica DM2000) to determine cannula placement, and only animals with bilateral correctly placed cannulas were included (for an example see **Figure 1A**).

**Figure 1.**
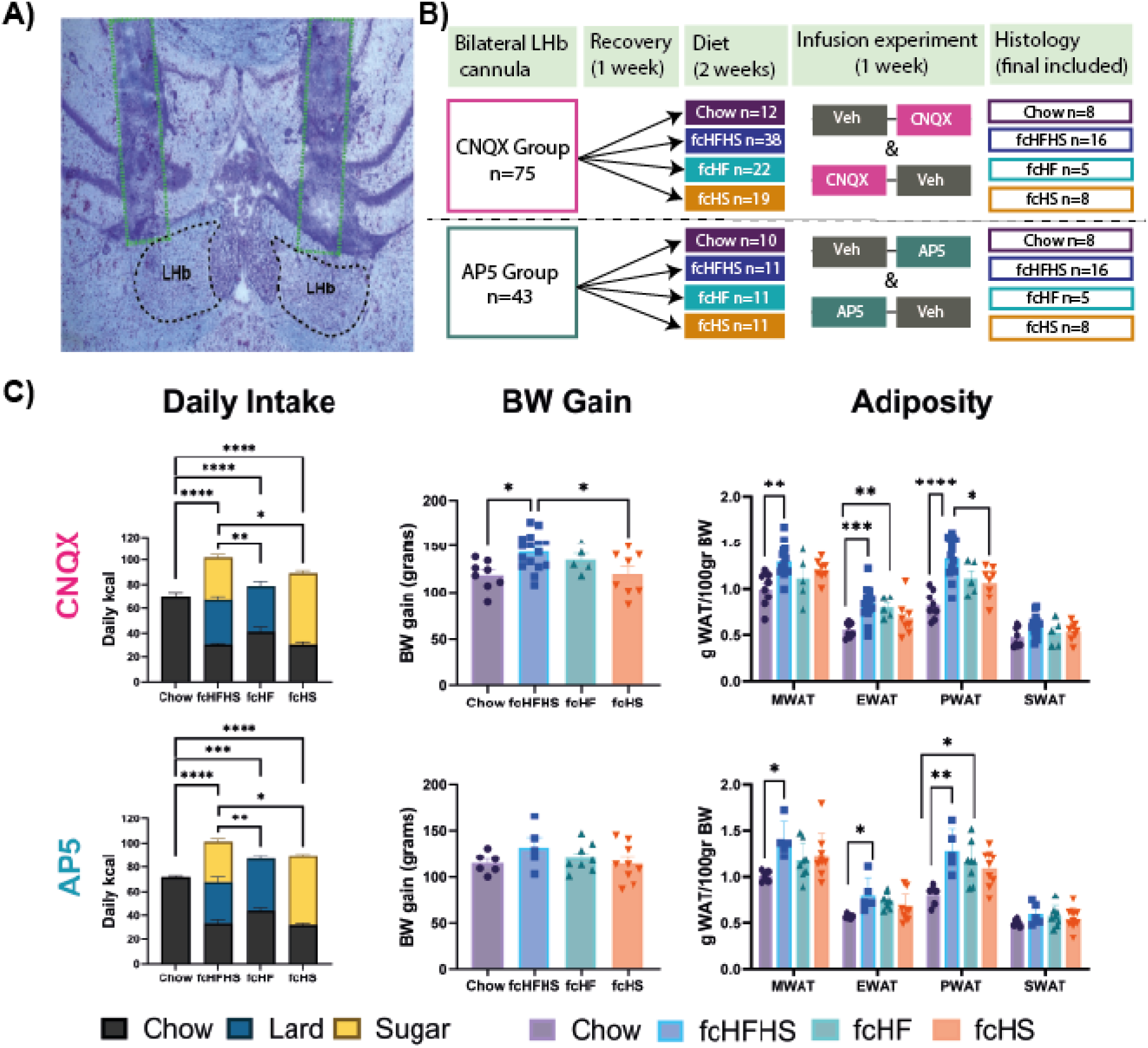
Experimental overview, daily intake, body weight gain, and adiposity per diet group in both experiments (CNQX and AP5). **A)** Example image from correct cannula placement. **B)** Timeline of the experiment with animal numbers. For the CNQX group, daily intake **(C)** of the various diet components is displayed, and body weight gain throughout the experiment. The fcHFHSD group gained significantly more body weight (in grams) compared to the chow diet group. In terms of adiposity, fcHFHSD animals had the greatest adiposity for the MWAT, EWAT, and PWAT. For SWAT, no differences between diet groups were observed. For the AP5 group, daily intake of the various diet components is displayed, and body weight gain throughout the experiment, with no differences between diet groups. In terms of adiposity, fcHFHS animals had the greatest adiposity for the MWAT, EWAT, and PWAT. For SWAT, no differences between diet groups were observed. For both CNQX and AP5 groups, total daily caloric intake was significantly higher in all diet groups compared to chow diet, and lower in the fcHFD and fcHSD groups compared to the fcHFHSD group. BW=bodyweight; WAT=white adipose tissue; MWAT=mesenteric WAT; EWAT=epididymal WAT; PWAT=perirenal WAT; SWAT=subcutaneous WAT. Data shown mean ± SEM. Sample sizes: CNQX: 8 (chow controls), 16 (fcHFHSD), 5 (fcHFD), 8 (fcHSD); AP5: 6 (chow controls), 5 (fcHFHSD), 8 (fcHFD), 9 (fcHSD). Statistics were performed with one-way ANOVAs. * = p<0.05 as tested with a posthoc Tukey’s test.

### Inclusion and exclusion

For the CNQX infusion experiment, a total of 75 rats were used, distributed over four diet groups (i.e. chow, fcHFHSD, fcHFD, and fcHSD). One CNQX-fcHFHSD-fed rat was removed from the experiment as he lost too much weight after infusion. One other fcHFHSD-fed animal was excluded due to too little fat consumption (<10% of average daily calories). Animals with misplaced cannulas (uni-or bilaterally) were excluded as well, leaving 16/38 fcHFHSD-fed animals with bilaterally correctly placed cannulas to be included. For the other diet groups, 5/22 fcHFD-fed animals, 8/19 fcHSD-fed animals, and 8/12 chow-fed animals were included based on cannula placement, leading to an overall inclusion of 37 rats.

For the AP5 infusion experiment, similar criteria were used. This resulted in 5/11 fcHFHSD-fed, 8/11 fcHFD-fed, 9/11 fcHSD-fed, and 6/10 chow-fed animals to be included. Across all diet groups, 28/43 rats were included for the AP5 infusions (**Figure 1B**).

### Statistical analysis

All data are expressed as the mean ± SEM. Graph Pad Prism (Version 9.1.0) software was used to visualize data and perform statistical analyses. To compare the effects of drug infusion (VEH vs. CNQX and AP5) on intake of individual diet components, two-way repeated measures analysis of variance (ANOVA), followed by post hoc analysis (Šidák), was used. For the acute time points (1h), paired t-tests were performed for each diet group. *P*-values less than 0.05 were considered to be statistically significant, and *P*-values below 0.1 were considered a trend.

## Results

### Body composition and daily caloric intake

In the CNQX infusion cohort, fcHFHSD, fcHFD, and fcHSD rats had greater daily caloric intake compared to Chow controls, whereas fcHFHSD rats had greater daily caloric intake compared to fcHFD and fcHSD rats **(Figure 1A)**. These changes in caloric intake were reflected in body weight gain throughout the experiment, which was increased in fcHFHSD rats compared to Chow controls **(Figure 1B)**. Furthermore, fcHFHSD rats had greater MWAT, EWAT, and PWAT, but not SWAT, mass compared to Chow controls (**Figure 1C**). In the AP5 infusion cohort, daily caloric intake was increased in fcHFHSD, fcHFD, and fcHSD rats compared to Chow controls **(Figure 1D**). The fcHFHSD rats had a significantly higher daily intake than the fcHFD and fcHSD rats **(Figure 1D**). This was not reflected in body weight gain over the course of the experiment as there were no differences between diet groups (**Figure 1E**). In terms of adiposity, the fcHFHSD rats had increased MWAT, EWAT, and PWAT, but not SWAT, mass compared to Chow controls, with no differences between fcHFD or fcHSD rats (**Figure 1F**).

### Glutamatergic inhibition of LHb does not acutely affect caloric intake

Infusion of an AMPA/kainate antagonist (CNQX) or NMDA receptor antagonist (AP5) into the LHb did not significantly modulate caloric intake one-hour following infusion in Chow, fcHFHSD, fcHFD, and fcHSD rats (**Figure 2**). One hour after intra-LHb CNQX infusion, total caloric intake was not altered in chow-, fcHFD-or fcHSD-fed animals. However, in the fcHFHSD-fed animals, a trend towards a decrease in total calories consumed was observed (**Figure 2A**). This was accompanied by no changes in intake of any of the specific components 1 hour after CNQX infusion (**Figure 2B)**. Similarly, infusion of AP5, an NMDA receptor antagonist, did not change total calories consumed after 1 hour in any of the diet groups, nor consumption of any diet component specifically (**Figure 2C-D**). These results suggest that glutamatergic inhibition, either by blocking NMDA or AMPA receptors in the LHb, does not acutely alter food intake.

**Figure 2.**
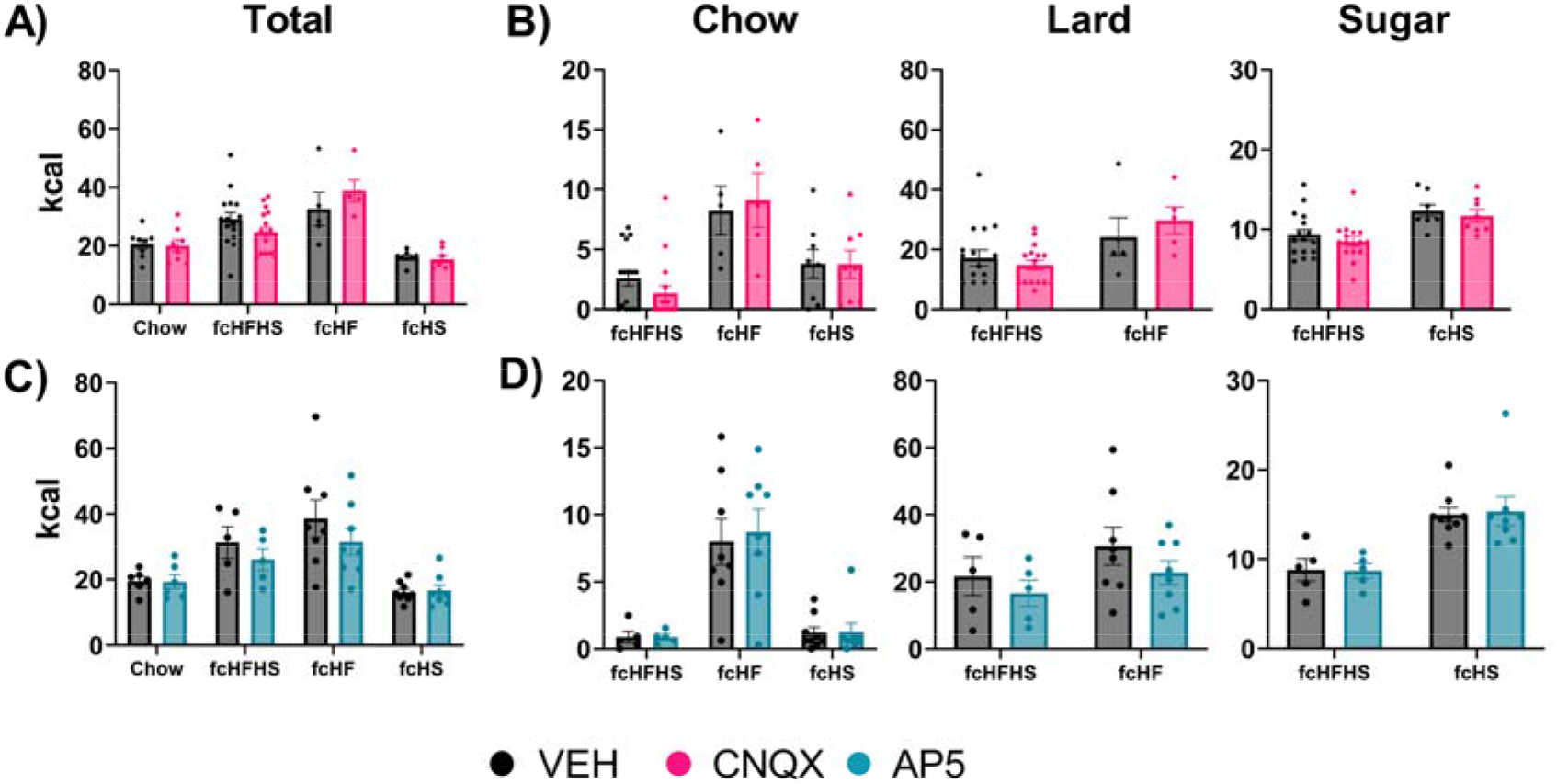
No acute effects (1 hour) on intake of all diet components in 4 different diet groups after CNQX or AP5 infusion in the LHb. **A)** Total caloric intake was not altered 1 hour after intra-LHb infusion of CNQX for chow-(t(7)=0.26, p=0.80), fcHFD-(t(4)=0.84, p=0.45), fcHSD-fed (t(7)=0.53, p=0.61) or fcHFHSD-fed (t(15)=1.80, p=0.09•) animals. **B)** Chow, lard an sugar intake for the fcHFHS, fcHF and fcHS-fed groups 1 hour after CNQX infusion was not altered (**fcHFHSD** chow t(15)=1.36, p=0.19; lard t(15)=1.10, p=0.29; sugar t(15)=1.35, p=0.19; **fcHFD** chow t(4)=0.29, p=0.78; lard t(4)=0.73, p=0.50; **fcHSD** chow t(7)=0.03, p=0.98; sugar t(7)=0.78, p= 0.46). **C)** Total caloric intake for all diet groups 1 hour after AP5 infusion was not altered (**chow** t(5)=0.10, p=0.92; **fcHFHSD** t(4)=1.48, p=0.21; **fcHFD** t(7)=1.43, p=0.19; **fcHSD** t(7)=0.35, p=0.73). **D)** Chow, lard an sugar intake for the fcHFHS, fcHF and fcHS-fed groups 1 hour after AP5 infusion was not altered(**fcHFHSD** chow t(4)=0.00, p=0.99; lard t(4)=1.53, p=0.20; sugar t(4)=0.17, p=0.87; **fcHFD** chow t(7)=0.52, p=0.62; lard t(7)=1.59, p=0.16; **fcHSD** chow t(7)=0.82, p=0.44 ; sugar t(7)=0.14, p=0.89). Sample sizes CNQX: 8 (chow), 16 (fcHFHS), 5 (fcHF), 8 (fcHS); AP5: 6 (chow), 5 (fcHFHS), 8 (fcHF), 9 (fcHS). Paired t-tests indicated no significant differences; data displayed mean ± SEM.

### NMDAR antagonism in the LHb induces a delayed decrease in lard intake in fcHFHSD rats

While there were no acute (1hr) effects of AP5 infusion, we were interested to see effects after a longer time period. Therefore, we included time points to measure food consumption after 10 and 24 hours (**Figure 3)**. Intra-LHb infusion of AP5 did not change total calorie consumption in any of the diet groups, however in the fcHFD-fed rats, total intake tended to decrease after AP5 infusion. In terms of specific components, this minor decrease in total calories seems to be driven by fat consumption in both fcHFHSD and fcHFD rats. Both chow and sugar consumption were not affected by intra-LHb AP5 infusion in any of the diet groups.

**Figure 3.**
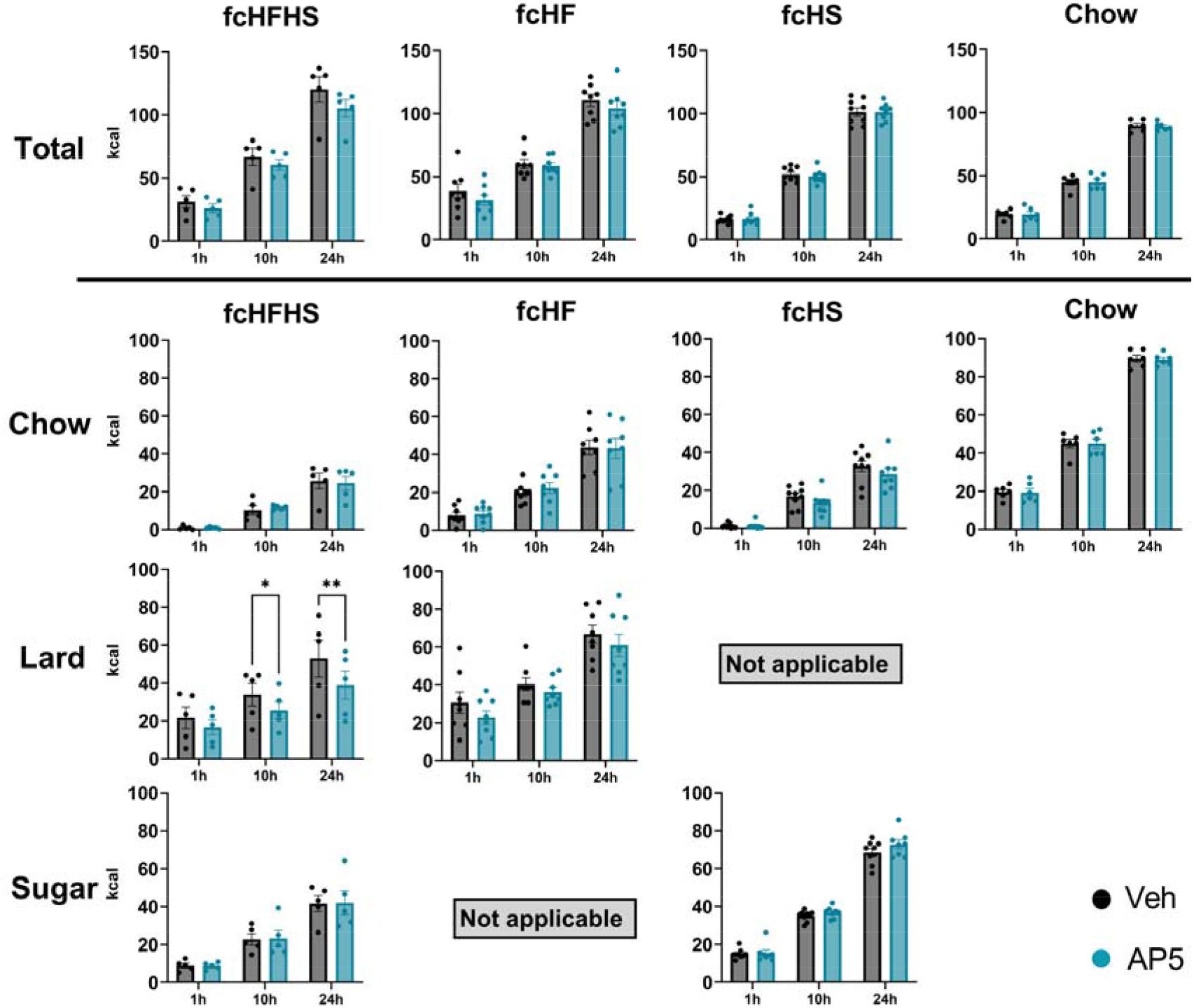
Effects on infusions of AP5 onto the LHb in different diet groups (chow/fcHFHS/fcHF/fcHS) on food components. Total caloric intake was not affected (**fcHFHSD** drug p=0.31, time x drug p=0.48; **fcHSD** drug p=0.96, time x drug p=0.75; **chow** drug p=0.88, time x drug p=0.98; **fcHFD** drug p=0.07^•^, time x drug p=0.65). Only in the fcHFHSD group, long-term effects on fat consumption were observed, after 10 hours and 24 hours (**fcHFHSD** drug p=0.06^•^, drug x time p=0.08^•^, posthoc 10h p=0.026*, 24h p=0.001**; **fcHFD** lard drug p=0.07^•^, drug x time p=0.79). For none of the other groups and time points, significant differences between consumption after vehicle and AP5 infusion were observed (**fcHFHSD** *chow* drug p=0.97, time x drug p=0.78; *sugar* drug p=0.94, time x drug p=0.98; **fcHFD** *chow* drug p=0.53, drug x time p=0.42; **fcHSD** *chow* drug p=0.16, drug x time p=0.09^•^; *sugar* drug =0.24, drug x time p=0.24). Sample sizes AP5: 6 (chow), 5 (fcHFHS), 8 (fcHF), 9 (fcHS). Two-way repeated measures ANOVA with Šidák post-hoc when interaction term showed a trend (p<0.1). Data displayed mean ± SEM.

### Effects of combined AMPA and NMDA antagonism in the LHb

Next, we focused on fcHFHSD-fed rats as there was a trend observed. We performed an experiment in a separate cohort of fcHFHSD rats to study the potential additive effects of a combined infusion of CNQX and AP5, and thus simultaneously block NMDA and AMPA receptors. In this cohort, we included time points at 1, 3, 6, and 24 hours to minimize the risk of missing the timing of a potential effect, and in addition also performed another CNQX infusion. When comparing CNQX and CNQX+AP5 infusion (in the same animals on fcHFHSD), total intake did not change at any of the time points (**Figure 4A, C)**. After CNQX infusion, no changes in intake of any of the specific components was observed (**Figure 4B**). Interestingly, when looking at specific components after CNQX+AP5 infusion, sugar intake was decreased at a later time point noted while other components did not change (**Figure 4D**).

**Figure 4.**
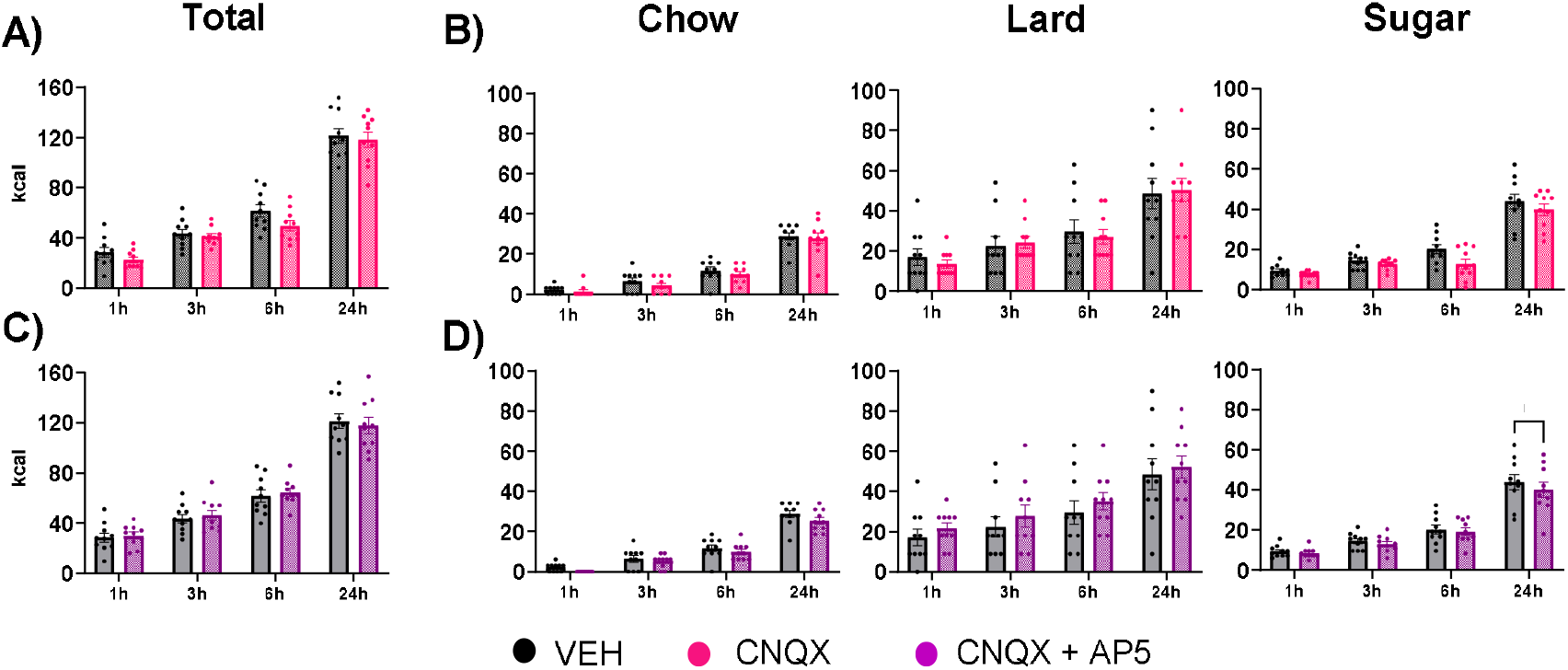
In fcHFHS-fed animals, infusion of CNQX combined with AP5 decreased sugar intake after 24 hours. **A)** Total calorie intake for all time points after CNQX infusion was not altered (drug p=0.24, time x drug p=0.39) **B)** Chow, lard, and sugar intake over time after CNQX infusion was not altered (chow drug p=0.39, time x drug p=0.93; lard drug p=0.89, time x drug p=0.51; sugar drug p=0.14, time x drug p=0.17). **C)** Total caloric intake for all time points after CNQX+AP5 infusion was not altered (drug p=0.83, time x drug p=0.32). **D)** Late sugar intake (24h) was decreased after CNQX+AP5 infusion (sugar drug p=0.04*, time x drug p=0.32, posthoc 24h p=0.02*), while not affecting chow and lard consumption (chow drug p=0.12, time x drug p=0.64; lard drug p=0.23; time x drug p=0.92). The sample size was 10 animals. Statistics were performed using Two-way repeated measures ANOVA with Šidák posthoc if necessary, data displayed mean ± SEM.

## Discussion

Here we demonstrated that pharmacological inhibition of LHb input did not acutely affect feeding in chow-, fcHFHSD-, fcHFD-, or fcHSD-fed rats, contrary to our hypothesis. Nevertheless, inhibition with the NMDA receptor antagonist AP5 tended to decrease fat intake at 10h and 24h following infusion, and this effect was more profound in the fcHFHSD-fed rats compared to the fcHFD-fed rats (fcHFHSD p=0.06; fcHFD p=0.07). This delayed effect on fat intake was not observed following infusion of the AMPA receptor antagonist CNQX, or following simultaneous infusion of AP5 and CNQX. However, after this combined infusion of CNQX and AP5, a delayed decrease in sugar intake was observed in a fcHFHSD-fed group at 24h following infusion.

While the observed effects on feeding were small and delayed, we did observe that inhibition of glutamatergic transmission in the LHb decreased palatable food intake, which is in contrast with several studies that showed that stimulation of specific glutamatergic inputs to the LHb decreased palatable food intake[29, 31]. Inhibition of LHb neurons will likely lower their indirect inhibitory effect on VTA dopaminergic neurons and thereby increase motivational drive for food seeking and/or consumption. In accordance with this, a recent study demonstrated that optogenetic inhibition of a glutamatergic LH -> LHb connection increased palatable liquid consumption [29]. Another recent report showed increased food intake after inhibition of glutamatergic NPY1R LHb neurons in mice maintained on an HFD [50]. Another study, where basal forebrain input to the LHb was activated, reported a decrease in food intake during the fasted state [31]. Collectively, these studies suggest that specific connections regulate consummatory behavior via the LHb in a specific manner. The pharmacological approach used in this study blocks glutamatergic transmission in the LHb in a general manner. Thus, it could be that an overall decrease in glutamatergic signaling in the LHb without a specific context (like stress) does not affect food intake but affects other behaviors such as anxiety-related behaviors, and introduces a more aversive state, whereas specific connections within different contexts are involved in the regulation of food intake. Indeed, while previously mentioned specific LHb inputs affect feeding, non-specific pharmacological inhibition of AMPA signaling in the LHb produced less exploratory behavior in a novel environment [64] and produced stress-like behavior in a water maze test as well as memory deficits [65]. Notably, the (small) effects on diet component intake that we did observe were exclusively in the groups that consumed a diet that included lard (*i*.*e*. fcHFHSD and fcHFD). LHb output modulates dopamine signaling, and previously we have shown that dopamine manipulations mostly impact consumption of lard, and rarely sugar intake [68]. This is in line with our observation that the observed effects on palatable diet component consumption seem specific for lard, and not present in fcHSD-fed rats.

We used a pharmacological approach with glutamatergic antagonists and thus effects depend on the number of available receptors to block at the moment of the infusion. For the NMDA antagonist, this receptor availability will depend on neuronal activity which, in the LHb, is known to display a day-night rhythm [12, 13], is increased during consumption of an HFD [50] or during a depressive-like state [40-42], and is sensitive to stress [50]. Specifically, the AMPA/NMDA receptor ratio in LHb synapses changes upon stress, turning the LHb signal more aversive and driving cognitive impairments [66, 67]. Taken together, these observations suggest that the main effect of pharmacological inhibition of the LHb is highly context-dependent, such as on which time of day and during which state animal infusions are performed. Given the sensitivity of the LHb to stress and its role in anxiety and depression, these factors could all acutely modulate LHb activity and its subsequent response to pharmacological blockade of glutamatergic input by altering the AMPA/NMDA ratio. To further our understanding of the LHb and its role in the consumption of palatable diets, it would be of interest to investigate if LHb AMPA/NMDA receptor ratios are affected during consumption of the various diet types used in this study (*i*.*e*. fcHFHSD, fcHFD, fcHSD) or various physiological states (*i*.*e*. stressed, anxious). As stress-induced overeating of a palatable diet was ablated when inhibiting NPY1R in the LHb [50], we would hypothesize that only during a stress-challenged state, LHb AMPA/NMDA receptor ratio is altered to modulate consumption of palatable diets.

Our study has a few limitations. First, various group sample sizes are small (n=5), which limits our statistical power. Secondly, we only used a single dose of each antagonist, and this dose was based on the literature [48, 63]. As we mostly observed trending effects, performing a dose-response curve for each antagonist might provide extra information if a higher concentration would have a more substantial effect. Third, we recorded consummatory behavior, but not locomotor activity or other measures of general behavior like grooming or rearing, and we only recorded consummatory behavior on specific, pre-determined time points (*i*.*e*. 1h, 10h, and 24h post-infusion). Using continuous and non-disruptive measurements of consummatory behavior could provide more detailed information on the role of the LHb during consumption of palatable diets. For future experiments, it is also important to include behavioral paradigms, such as the elevated plus maze or conditioned place aversion, as a positive control for the blockade of glutamatergic LHb input. We conclude that LHb AMPA and NMDA signaling play a minor role in the modulation of palatable food intake, in a delayed manner, using our experimental settings.

## Acknowledgements

This work was supported by an AMC PhD fellowship grant awarded by the AMC Executive Board.

## Author Contributions

MS, JDM, and SElF designed the research with input from AASvI, TK, and LE. MS conducted the research with assistance from MTS, JLvB, AASvI, TK, and LE. MS performed the analysis and histology with assistance of MTS and JLvB. MS, JDM, and SElF wrote the manuscript. All authors read and approved the manuscript.

## References

1. Bianco, I.H. and S.W. Wilson, The habenular nuclei: a conserved asymmetric relay station in the vertebrate brain. Philosophical Transactions of the Royal Society B: Biological Sciences, 2009. 364(1519): p. 1005–1020.

2. Agetsuma, M., et al., The habenula is crucial for experience-dependent modification of fear responses in zebrafish. Nature neuroscience, 2010. 13(11): p. 1354–1356.

3. Stephenson-Jones, M., et al., Evolutionary conservation of the habenular nuclei and their circuitry controlling the dopamine and 5-hydroxytryptophan (5-HT) systems. Proceedings of the National Academy of Sciences, 2012. 109(3): p. E164–E173.

4. Namboodiri, V.M.K., J. Rodriguez-Romaguera, and G.D. Stuber, The habenula. Current Biology, 2016. 26(19): p. R873–R877.

5. Marburg, O., The structure and fiber connections of the human habenula. Journal of Comparative Neurology, 1944. 80(2): p. 211–233.

6. Aizawa, H., et al., Molecular characterization of the subnuclei in rat habenula. Journal of Comparative Neurology, 2012. 520(18): p. 4051–4066.

7. Geisler, S., K.H. Andres, and R.W. Veh, Morphologic and cytochemical criteria for the identification and delineation of individual subnuclei within the lateral habenular complex of the rat. Journal of Comparative Neurology, 2003. 458(1): p. 78–97.

8. Wagner, F., T. Stroh, and R.W. Veh, Correlating habenular subnuclei in rat and mouse by using topographic, morphological, and cytochemical criteria. Journal of Comparative Neurology, 2014. 522(11): p. 2650–2662.

9. Weiss, T. and R. Veh, Morphological and electrophysiological characteristics of neurons within identified subnuclei of the lateral habenula in rat brain slices. Neuroscience, 2011. 172: p. 74–93.

10. Andres, K.H., M.V. Düring, and R.W. Veh, Subnuclear organization of the rat habenular complexes. Journal of Comparative Neurology, 1999. 407(1): p. 130–150.

11. Lecourtier, L. and P.H. Kelly, A conductor hidden in the orchestra? Role of the habenular complex in monoamine transmission and cognition. Neuroscience & Biobehavioral Reviews, 2007. 31(5): p. 658–672.

12. Zhao, H. and B. Rusak, Circadian firing-rate rhythms and light responses of rat habenular nucleus neurons in vivo and in vitro. Neuroscience, 2005. 132(2): p. 519–528.

13. Guilding, C., A. Hughes, and H. Piggins, Circadian oscillators in the epithalamus. Neuroscience, 2010. 169(4): p. 1630–1639.

14. Baltazar, R.M., L.M. Coolen, and I.C. Webb, Diurnal rhythms in neural activation in the mesolimbic reward system: critical role of the medial prefrontal cortex. European Journal of Neuroscience, 2013. 38(2): p. 2319–2327.

15. Hikosaka, O., et al., Habenula: crossroad between the basal ganglia and the limbic system. Journal of Neuroscience, 2008. 28(46): p. 11825–11829.

16. Sutherland, R.J.J.N. and B. Reviews, The dorsal diencephalic conduction system: a review of the anatomy and functions of the habenular complex. 1982. 6(1): p. 1–13.

17. Brown, P.L. and P.D. Shepard, Functional evidence for a direct excitatory projection from the lateral habenula to the ventral tegmental area in the rat. Journal of neurophysiology, 2016. 116(3): p. 1161–1174.

18. Lammel, S., et al., Input-specific control of reward and aversion in the ventral tegmental area. Nature, 2012. 491(7423): p. 212–217.

19. Jhou, T.C., et al., The mesopontine rostromedial tegmental nucleus: a structure targeted by the lateral habenula that projects to the ventral tegmental area of Tsai and substantia nigra compacta. Journal of Comparative Neurology, 2009. 513(6): p. 566–596.

20. Hong, S., et al., Negative reward signals from the lateral habenula to dopamine neurons are mediated by rostromedial tegmental nucleus in primates. Journal of Neuroscience, 2011. 31(32): p. 11457–11471.

21. Stamatakis, A.M. and G.D. Stuber, Activation of lateral habenula inputs to the ventral midbrain promotes behavioral avoidance. Nature neuroscience, 2012. 15(8): p. 1105.

22. Herkenham, M. and W.J. Nauta, Efferent connections of the habenular nuclei in the rat. Journal of Comparative Neurology, 1979. 187(1): p. 19–47.

23. Hong, S. and O. Hikosaka, The globus pallidus sends reward-related signals to the lateral habenula. Neuron, 2008. 60(4): p. 720–729.

24. Shabel, S.J., et al., Input to the lateral habenula from the basal ganglia is excitatory, aversive, and suppressed by serotonin. Neuron, 2012. 74(3): p. 475–481.

25. Wallace, M.L., et al., Genetically distinct parallel pathways in the entopeduncular nucleus for limbic and sensorimotor output of the basal ganglia. Neuron, 2017. 94(1): p. 138–152. e5.

26. Li, H., D. Pullmann, and T.C. Jhou, Valence-encoding in the lateral habenula arises from the entopeduncular region. Elife, 2019. 8: p. e41223.

27. Lazaridis, I., et al., A hypothalamus-habenula circuit controls aversion. Molecular psychiatry, 2019: p. 1.

28. Lecca, S., et al., Aversive stimuli drive hypothalamus-to-habenula excitation to promote escape behavior. Elife, 2017. 6: p. e30697.

29. Stamatakis, A.M., et al., Lateral hypothalamic area glutamatergic neurons and their projections to the lateral habenula regulate feeding and reward. Journal of Neuroscience, 2016. 36(2): p. 302–311.

30. Golden, S.A., et al., Basal forebrain projections to the lateral habenula modulate aggression reward. Nature, 2016. 534(7609): p. 688–692.

31. Swanson, J.L., et al., Activation of basal forebrain-to-lateral habenula circuitry drives reflexive aversion and suppresses feeding behavior. Scientific Reports, 2022. 12(1): p. 22044.

32. Root, D.H., et al., Role of glutamatergic projections from ventral tegmental area to lateral habenula in aversive conditioning. Journal of Neuroscience, 2014. 34(42): p. 13906–13910.

33. Stamatakis, A.M., et al., A unique population of ventral tegmental area neurons inhibits the lateral habenula to promote reward. Neuron, 2013. 80(4): p. 1039–1053.

34. Shabel, S.J., et al., GABA/glutamate co-release controls habenula output and is modified by antidepressant treatment. Science, 2014. 345(6203): p. 1494–1498.

35. Meye, F.J., et al., Shifted pallidal co-release of GABA and glutamate in habenula drives cocaine withdrawal and relapse. Nature neuroscience, 2016. 19(8): p. 1019–1024.

36. Barker, D.J., et al., Lateral preoptic control of the lateral habenula through convergent glutamate and GABA transmission. Cell reports, 2017. 21(7): p. 1757–1769.

37. Nuno-Perez, A., et al., Lateral habenula gone awry in depression: bridging cellular adaptations with therapeutics. Frontiers in neuroscience, 2018. 12: p. 485.

38. Proulx, C.D., O. Hikosaka, and R. Malinow, Reward processing by the lateral habenula in normal and depressive behaviors. Nature neuroscience, 2014. 17(9): p. 1146.

39. Yang, Y., et al., Lateral habenula in the pathophysiology of depression. Current opinion in neurobiology, 2018. 48: p. 90–96.

40. Caldecott-Hazard, S., J. Mazziotta, and M. Phelps, Cerebral correlates of depressed behavior in rats, visualized using 14C-2-deoxyglucose autoradiography. Journal of Neuroscience, 1988. 8(6): p. 1951–1961.

41. Mirrione, M.M., et al., Increased metabolic activity in the septum and habenula during stress is linked to subsequent expression of learned helplessness behavior. Frontiers in human neuroscience, 2014. 8: p. 29.

42. Andalman, A.S., et al., Neuronal dynamics regulating brain and behavioral state transitions. Cell, 2019. 177(4): p. 970–985. e20.

43. Morris, J., et al., Covariation of activity in habenula and dorsal raphe nuclei following tryptophan depletion. Neuroimage, 1999. 10(2): p. 163–172.

44. Strotmann, B., et al., Mapping of the internal structure of human habenula with ex vivo MRI at 7T. Frontiers in human neuroscience, 2013. 7: p. 878.

45. Schmidt, F.M., et al., Habenula volume increases with disease severity in unmedicated major depressive disorder as revealed by 7T MRI. European archives of psychiatry and clinical neuroscience, 2017. 267(2): p. 107–115.

46. Ely, B.A., et al., Resting-state functional connectivity of the human habenula in healthy individuals: Associations with subclinical depression. Human brain mapping, 2016. 37(7): p. 2369–2384.

47. Meye, F.J., et al., Cocaine-evoked negative symptoms require AMPA receptor trafficking in the lateral habenula. Nature neuroscience, 2015. 18(3): p. 376–378.

48. Yang, Y., et al., Ketamine blocks bursting in the lateral habenula to rapidly relieve depression. Nature, 2018. 554(7692): p. 317.

49. Li, J., et al., Inhibition of AMPA receptor and CaMKII activity in the lateral habenula reduces depressive-like behavior and alcohol intake in rats. Neuropharmacology, 2017. 126: p. 108–120.

50. Ip, C.K., et al., Critical Role of Lateral Habenula Circuits in the Control of Stress-Induced Palatable Food Consumption. Neuron, 2023. 111: p. 1–18.

51. Fu, O., et al., Hypothalamic neuronal circuits regulating hunger-induced taste modification. Nature communications, 2019. 10(1): p. 1–14.

52. Rossi, M.A., et al., Transcriptional and functional divergence in lateral hypothalamic glutamate neurons projecting to the lateral habenula and ventral tegmental area. Neuron, 2021. 109(23): p. 3823–3837. e6.

53. Jhou, T.C. and P.J. Vento, Bidirectional regulation of reward, punishment, and arousal by dopamine, the lateral habenula and the rostromedial tegmentum (RMTg). Current Opinion in Behavioral Sciences, 2019. 26: p. 90–96.

54. Carlson, H.N., B.A. Christensen, and W.E. Pratt, Stimulation of mu opioid, but not GABAergic, receptors of the lateral habenula alters free feeding in rats. Neuroscience Letters, 2022. 771: p. 136417.

55. Companion, M.A., et al., Lateral habenula-projecting central amygdala circuits expressing GABA and NPY Y1 receptor modulate binge-like ethanol intake in mice. Addiction neuroscience, 2022. 3: p. 100019.

56. Slomp, M., et al., Stressing the importance of choice: Validity of a preclinical free-choice high-caloric diet paradigm to model behavioural, physiological and molecular adaptations during human diet-induced obesity and metabolic dysfunction. Journal of Neuroendocrinology, 2019. 31(5): p. e12718.

57. La Fleur, S., et al., A reciprocal interaction between food-motivated behavior and diet-induced obesity. International journal of obesity, 2007. 31(8): p. 1286–1294.

58. La Fleur, S., et al., A free-choice high-fat high-sugar diet induces changes in arcuate neuropeptide expression that support hyperphagia. International journal of obesity, 2010. 34(3): p. 537–546.

59. DiFeliceantonio, A.G., et al., Supra-additive effects of combining fat and carbohydrate on food reward. Cell metabolism, 2018. 28(1): p. 33–44. e3.

60. Small, D.M. and A.G. DiFeliceantonio, Processed foods and food reward. Science, 2019. 363(6425): p. 346–347.

61. de Araujo, I.E., M. Schatzker, and D.M. Small, Rethinking food reward. Annual review of psychology, 2020. 71: p. 139–164.

62. Perszyk, E.E., et al., Fat and carbohydrate interact to potentiate food reward in healthy weight but not in overweight or obesity. Nutrients, 2021. 13(4): p. 1203.

63. Velazquez, A.S.B., Palatable food, clock genes and the reward circuitry. 2018, Université de Strasbourg; Universiteit van Amsterdam.

64. Lecourtier, L., L. Durieux, and V. Mathis, Alteration of Lateral Habenula Function Prevents the Proper Exploration of a Novel Environment. Neuroscience, 2023. 514: p. 56–66.

65. Mathis, V., et al., Excitatory transmission to the lateral habenula is critical for encoding and retrieval of spatial memory. Neuropsychopharmacology, 2015. 40(12): p. 2843–2851.

66. Shabel, S.J., et al., Stress transforms lateral habenula reward responses into punishment signals. Proceedings of the National Academy of Sciences, 2019. 116(25): p. 12488–12493.

67. Nuno-Perez, A., et al., Stress undermines reward-guided cognitive performance through synaptic depression in the lateral habenula. Neuron, 2021. 109(6): p. 947–956. e5.

68. Joshi, A., et al., Dopamine D1 receptor signalling in the lateral shell of the nucleus accumbens controls dietary fat intake in male rats. Appetite, 2021. 167: p. 105597.

